# Keystone taxa responsible for the microbial community stability and performance of activated sludges

**DOI:** 10.1101/2023.02.26.530128

**Authors:** Xiaonan Liu, Miaoxiao Wang, Bingwen Liu, Xiaoli Chen, Liyun An, Yong Nie, Xiao-Lei Wu

## Abstract

**Background:** The functions and stability of a community depend on its species, which form complex interaction networks. The keystone taxa identified by network analysis are generally considered to play a vital role in the structure and function of microbial communities, but there is no uniformly accepted operational definition of such taxa. Further, what species and how they affect the community’s stability and function are still poorly understood.

**Methods:** To solve this problem, we performed a large-scale network analysis of the microbial communities residing in 1186 activated sludge (AS) samples.

**Results:** We found that the AS co-occurrence network is a typical scale-free network. While most taxa in the AS co-occurrence network have little association, there are still a small number of taxa that are strongly interconnected. We defined a group of keystone taxa that have an important impact on network stability. Further analysis results indicate that the communities harboring the keystone taxa maintain higher stability, but these communities possess lower pollutant removal rates. In addition, we found that keystone taxa were more likely to appear in samples with lower sludge load.

**Conclusions:** Our work identified the keystone taxa that maintain the stability of microbial communities in the AS systems but at the cost of reducing their function. This finding shed light on the relationship between composition, stability, and function within microbial communities. It also provides novel insights into manipulating the function of microbial communities by modifying their composition.

## Background

Ecological stability is essential for the maintenance of biodiversity and therefore the sustainability of ecological functions and services(1). Species in ecosystems form complex interaction networks, in which gains and losses of species affect ecological stability. What and how species affect ecological network structure is a fundamental question in ecology(2). However, the answer to this question is poorly understood.

The traditional community stability measurement is limited by the long duration of the disturbance(3) and the difficulty in determining the cross disturbance. Recently, more and more methods to evaluate community stability have been proposed(4, 5), in which network analysis has been widely used by ecologists to explore interactions and community stability (5, 6). In ecological networks, complex systems can be simplified by taking species as nodes and their interactions as edges(7). The effects of species on the structure and stability of the community can be assessed by removing or replacing the node (species) in networks and measuring the topological properties of the perturbed networks(8). The species that are important for community structure and integrity of the community are usually called keystone taxa(9). Previous work on the effect of species on network stability has shown that removing the most connected species leads to more secondary extinctions than random species removal(10). In some cases, however, the removal of low-connected species can sometimes have a large effect on the community(10). These findings suggest that the keystone species that determine community stability is not strictly defined by high connectivity.

As one of the most important ecological participants, microorganisms drive the global biogeochemical processes and shape our planet(11, 12). The functional structure of microbial communities is highly variable, with functional traits often reflecting the environment in which the community is found(13). The relevance of microbial composition, community stability, and ecological function is of considerable interest. Though the significance of biological diversity for community stability and ecosystem function is highly debated(14), greater species diversity is generally associated with improved function and stability of microbial communities (15, 16). Previous studies have proposed a variety of explanations for the impact of diversity on ecosystem functions, including more efficient utilization of resources due to increased niche differentiation and resource exploitation(17), and the more likely emergence of species with crucial functional characteristics(18). The key to the difficulty in unifying these explanations is that the relationship between different species composition and community function and stability is inconsistent, which depends on the species’ function, abundance, and interaction with other species(19, 20). Recent studies have shown that there are also keystone taxa in microbial communities, which drive microbiome structure and function regardless of their abundance(21, 22). The identification of keystone taxa may help us analyze the impact of species on communities, regardless of their specific functions and species types.

Network analysis is thought to be a powerful tool for inferring keystone taxa from ecological communities(23, 24). Unlike the direct observation of predatory or competitive interactions in macro-ecological networks, the interactions between microorganisms can’t be directly observed in many cases. The co-occurrence network based on the statistical correlation of sequencing data can help identify potential biotic interactions in microbial communities(25, 26). As sequencing analyses of microbial communities become increasingly available, we will be better able to quantify complex interactions among community members(27). Based on the microbial co-occurrence network, a series of definitions of keystone taxa have emerged, including the highly associated species nodes called hubs taxa in some studies(6, 28), and the species nodes with high betweenness centrality(23, 29). There is no uniformly accepted operational definition of keystone taxa in ecology yet, especially in microbial ecology. Although a review suggested that the comprehensive scores of high mean degree, high closeness centrality, and low betweenness centrality should be used to determine the keystone taxa(30), the rationality of this definition has not been verified. Further, there is no unified understanding of how keystone taxa affect the community stability and functions.

The activated sludge (AS) of the wastewater treatment plant (WWTP) is the largest artificial microbial ecosystem with high biodiversity and biomass(31), and it can be controlled by various operating conditions(32). As a model ecosystem, AS has a great attraction in the analysis of microbial community structure and the exploration of ecological theory. In addition, an investigation from global WWTP provides an ideal dataset for comprehensively analyzing the microbial structure and function of AS system(31).

In this work, we chose AS system to explore the influence of keystone taxa on community stability and function. Firstly, we analyzed the microbial co-occurrence pattern of AS systems from global wastewater treatment plants. Then, we identified the keystone taxa in the AS network and explored its role in maintaining community stability and promoting wastewater treatment performance. These analyses help to reveal the impact of keystone taxa on community functions and AS system stability, and deepen our understanding of ecological theory.

## Results

### Microbial co-occurrence network of global AS systems

To obtain an interaction network of microorganisms in global AS systems, we constructed the microbial co-occurrence network of AS system, which contains 4992 nodes and 65457 edges (Fig. 1). These nodes are mainly composed of Proteobacteria, Bacteroidota, Myxococota, and Chloroflexi (Fig. 1a). Besides, microbial node taxa in the co-occurrence networks have the functions of nitrification, denitrification, phosphorus accumulation, glycogen accumulation, sulfate reduction, sulfur oxidation, and bulking and foaming, despite 73.08% of ASVs nodes have unassigned functions in the MiDAS 4 database (Fig. 1c). In addition, 19.49% of edges belonged to intra-phylum co-occurrence and 80.51% belonged to inter-phylum co-occurrence (Fig. 1d), suggesting that inter-phylum interactions dominate the interaction networks of AS system. Furthermore, the average degree of nodes (node degree is the number of connections to a node) of Acidobacteriota was significantly higher than that of other phyla, indicating that ASVs in Acidobacteriota were more prone to interactions (Fig. S1).

**Fig. 1.**
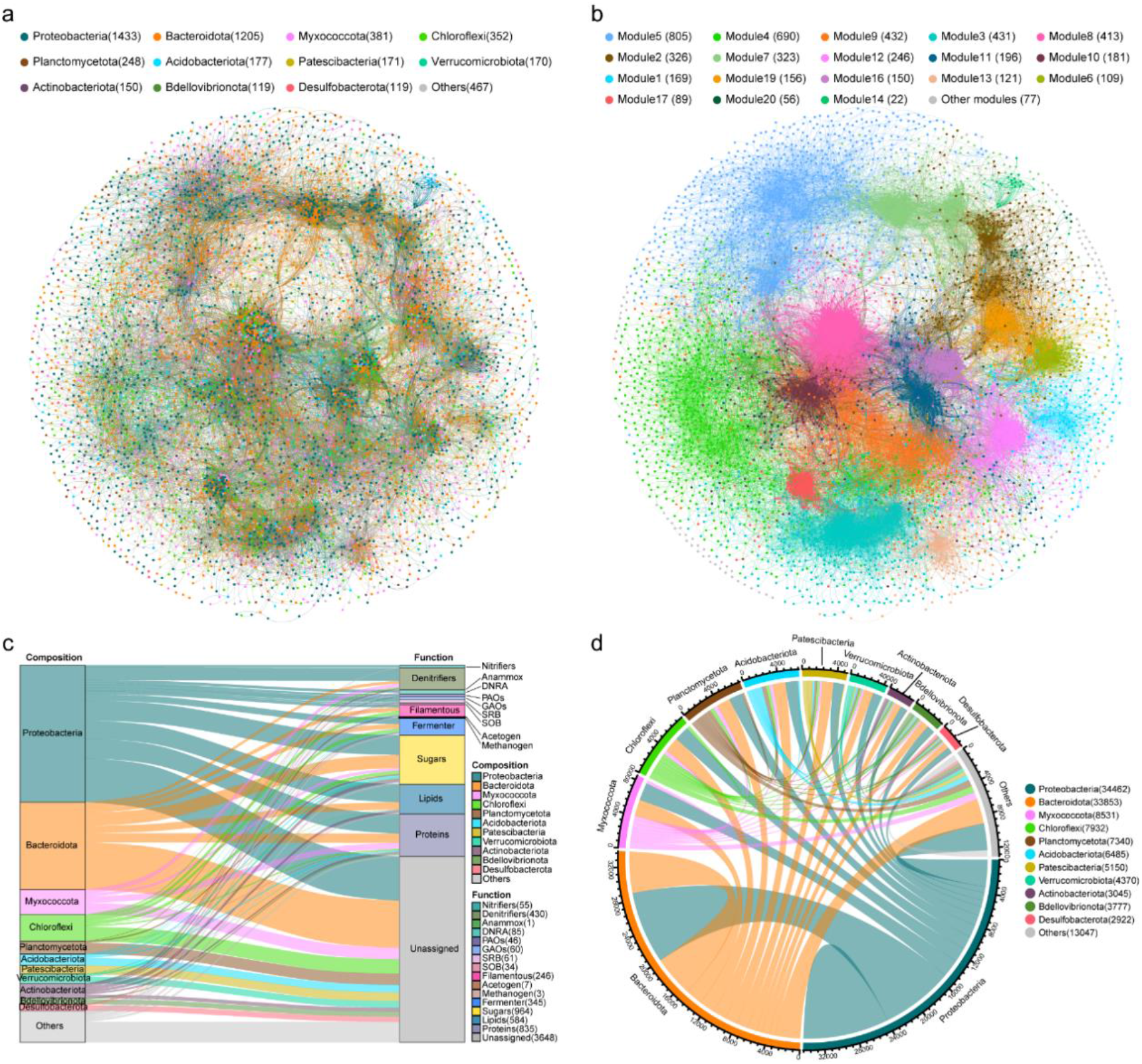
Microbial co-occurrence pattern of AS system. **(a)** AS co-occurrence network colored by phylum. **(b)** AS co-occurrence network colored by modularity class. **(c)** The composition and function annotation of nodes in the AS co-occurrence network. **(d)** Co-occurrence pattern of different phyla in AS system.

Topological characteristics of the co-occurrence network help to understand the interaction pattern between ASV nodes. Firstly, the standardization effect size (SES) was calculated to quantify the overall co-occurrence property of the ecosystem. The result showed that the SES of AS system was 2.84 (>2; Table 1), which implied the nonrandom community assembly patterns in AS system. Further analyzing the distribution of degrees of AS co-occurrence network, we found degrees of this network followed a power-law distribution (Fig. S2a), which was quite different from the Erdös - Réyni (ER) random networks with identical sizes (the same number of nodes and edges) (Fig. S2b). This result indicates that the AS network is a scale-free network, in which connections between its nodes are seriously uneven, specifically, a few ASVs nodes in the network have extremely many connections, while most ASVs nodes have only a few connections. Then, by comparing the topological properties of AS co-occurrence network and ER random networks with equal size (Table 1), we found that nodes in AS network are more highly connected than in identically sized random networks, as its higher clustering coefficient (CC) and small-world coefficient (σ=64.66)(33). In addition, the AS network had a modular structure, as the modularity (MD) was 0.8(>0.4)(34), and there were interactions among modules (Fig. 1b), suggesting the modularity characteristics in the assembly of community construction.

**Table 1.**
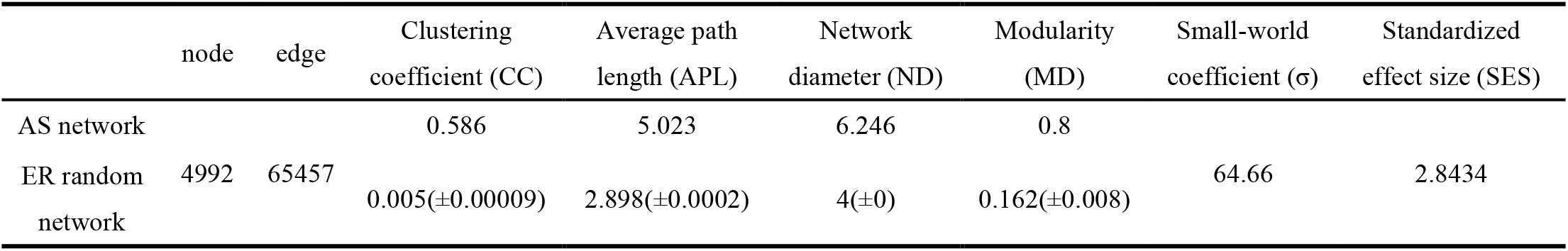
Topological properties of AS network and ER random network.

The stability of the network can reflect community stability in the face of environmental fluctuations to a certain extent. To analyze the stability(5) of AS network, we simulated and calculated the impact of species extinction on natural connectivity. The results showed that when species extinction was less than 56% of the community, the natural connectivity of the AS network was higher than that of the ER random network (Fig. S3), indicating that the AS network was robust, and non-random interactions of microorganisms in AS system was very important to maintain the stability of the community structure.

### Identification of keystone taxa sustaining the community stability of AS system

Then, we set out to identify the keystone taxa from the AS network and analyze their basic properties. Following the previous works (30, 35), we used node degree and betweenness centrality (meaning the number of shortest paths through the node in a network) as two key indexes. We defined the nodes with high degree (>100) and low betweenness centrality (<5000) in AS co-occurrence network as keystone taxa (Fig. 2a).

**Fig. 2.**
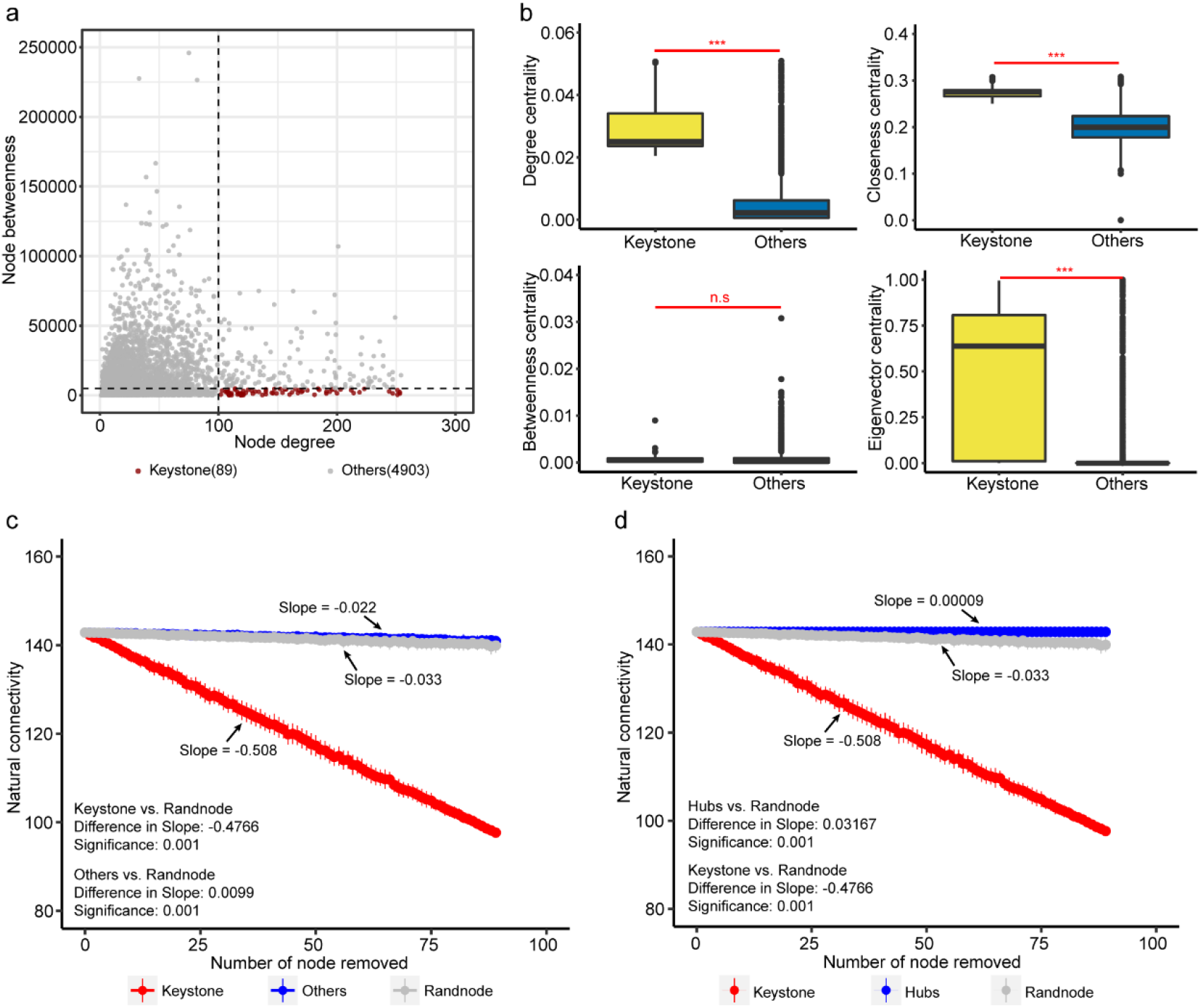
Identification and verification of keystone taxa. **(a)** Node degree and betweenness properties of keystone taxa. **(b)** Comparison of degree centrality, betweenness centrality, closeness centrality, and eigenvector centrality between keystone taxa and other taxa. Statistical analysis was performed using a two-sample Student’s t-test: ***, p < 0.001; n.s, p > 0.05, no significance. **(c)** The impact of randomly deleting keystone taxa or other taxa on the natural connectivity of AS network. (**d**) The impact of randomly deleting keystone taxa or hubs taxa on the natural connectivity of AS network. The statistical comparison of slopes was realized by using the *diffslope* function in the simba package of R 4.0.3.

To investigate whether these keystone taxa have a crucial impact on the structure of AS network as our expectation, we compared the node centrality of keystone taxa with others taxa. Node centrality quantifies the impact of each node and is thus usually used to measure the importance of nodes in the network(5, 36). The results showed that three indexes of node centrality of keystone taxa, including the degree centrality, closeness centrality, and eigenvector centrality, were all significantly higher than those of other taxa, while we observed no significant difference between the betweenness centrality of keystone and other taxa (Fig. 2b). This result implied that keystone taxa were of high importance in the AS network and may play an important role in maintaining the community structure of AS system.

Considering the high node importance of keystone taxa, we hypothesized that keystone taxa may play an important role in maintaining the stability of AS network. To test this hypothesis, we simulated and compared the impact of species extinction of keystone taxa and other taxa on the natural connectivity of the AS network. Natural connectivity is generally considered to reflect the stability of the network(5, 37). The simulation results showed that the influence of removing the keystone taxa on the natural connectivity of AS network was significantly higher than that of removing random species, while the influence of removing other taxa was significantly lower than that of removing random species (Fig. 2c). The significant impact of keystone taxa on the natural connectivity of AS network support our hypothesis that keystone taxa were crucial for maintaining the stability of AS network and may play an important role in maintaining the stability of AS microbial community.

Several previous studies applied another concept to define keystone taxa, namely hubs of a network are also often proposed as keystone taxa(6, 38, 39). Network hubs are a group of nodes with high connectivity within or among modules in the microbial co-occurrence network based on Zi and Pi indexes(28). We follow this definition, to identify 224 hub ASVs from the AS network (Fig. S4). However, the comparison of node importance between hubs taxa and the above-mentioned keystone taxa in AS network showed that node centralities of hubs taxa were significantly lower than those of keystone taxa (Fig. S5), implying that the keystone taxa identified based on our definition may have a greater impact on the structure of AS network than the hubs taxa. Furthermore, the deletion of hubs taxa had little impact on the natural connectivity of AS network, significantly lower than that of keystone taxa (Fig. 2d), which again confirms the rationality of the definition of keystone taxa in this paper.

We found that 40 of 89 keystone ASVs belonged to Proteobacteria and Bacteroidota, and about 10% belonged to Acidobacteriota (Fig. S6a), which is consistent with our previous finding that Acidobacteriota is more prone to interactions (Fig. S1). In addition, despite the small proportion of functional groups, keystone taxa still had the relevant functions of the chemical cycle of all substances including carbon, nitrogen, phosphorus, and sulfur (Fig. S6b). Furthermore, we found that the occurrence frequency (Fig. S6c) and average relative abundance (Fig. S6d) of keystone taxa were significantly lower than those of other taxa, indicating that keystone taxa are usually rare taxa possessing low abundance. In addition, we compared the contributions of keystone taxa with high-abundance and high-frequency core taxa in maintaining the stability of AS network. The results showed that the deletion of core taxa had little impact on the natural connectivity of AS network, significantly lower than that of keystone taxa (Fig. S7), indicating that keystone taxa had a higher impact on the stability of AS network than core taxa. In summary, our results demonstrated that the keystone species were independent of the species traits, such as their function and taxonomic distribution, but tend to have low occurrence frequency and abundance.

### Keystone taxa favor the community stability in AS systems

We further hypothesized that communities harboring keystone taxa are more stable than those communities not containing keystone taxa. Accordingly, we divided all samples into two groups according to whether they contained keystone ASVs, which we termed keystone samples (n=524) and no keystone samples (n=662) thereafter. The stability test of these two sample subnetworks with or without keystone taxa showed that the natural connectivity of the keystone samples subnetwork was significantly higher than that of no keystone group when the proportion of extinct species in the simulations was less than 60% (Fig. 3a). This result indicates that the presence of keystone taxa improves the stability of AS network. In addition, we found that the keystone samples subnetwork possesses a higher average degree and higher clustering coefficient of the subnetwork of keystone samples than those of the no keystone samples subnetwork (Table S1). This finding suggests that the presence of keystone taxa also increases the complexity of AS network, supporting the central ecological belief that complexity begets stability(6).

**Fig. 3.**
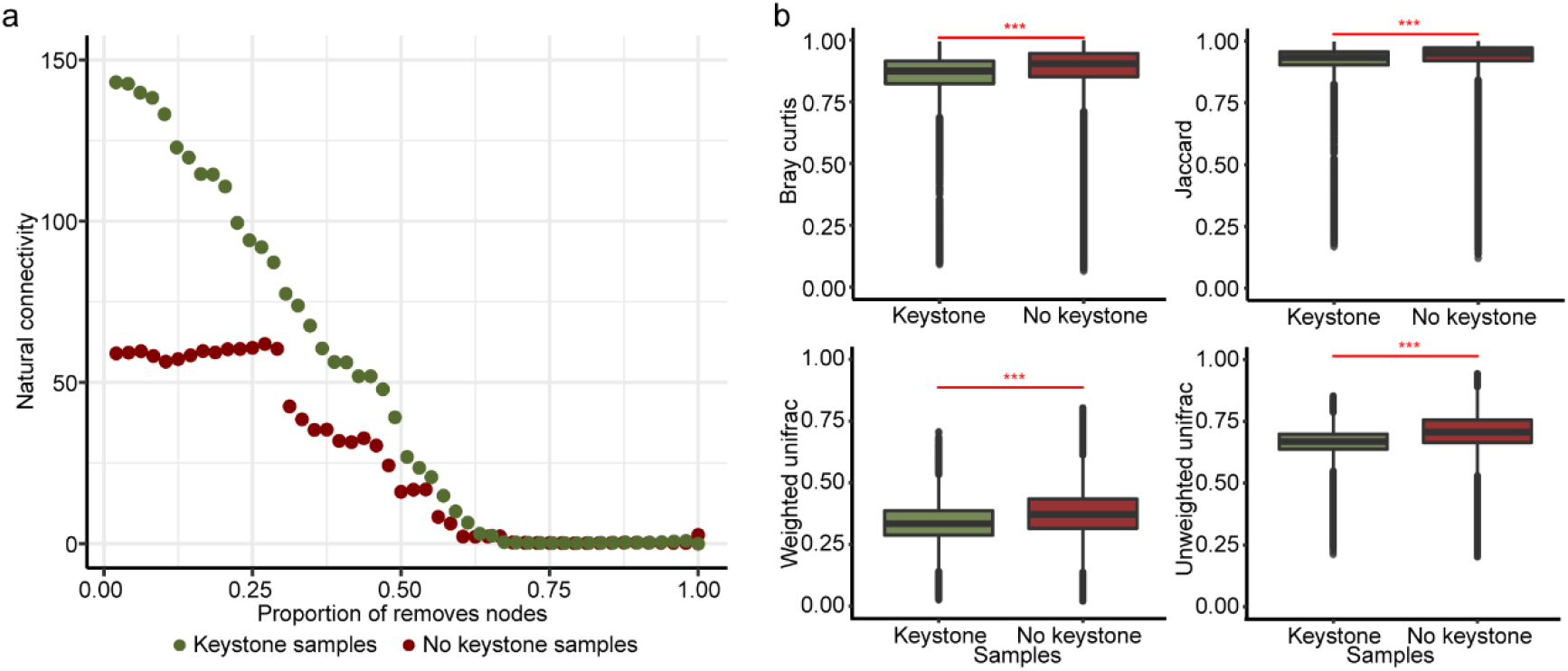
Contribution of keystone taxa to the community stability of AS system. **(a)** Comparison of the stability of subnetworks between keystone samples and no keystone samples. **(b)** Comparison of community dissimilarity between keystone samples and no keystone samples, measured by Bray-Curtis, Jaccard, weighted unifrac, and unweighted unifrac distance. Statistical analysis was performed using a two-sample Student’s t-test: ***, p < 0.001.

In addition, we compared the differences in community structure between keystone and no keystone samples. The results of principal coordinate analysis (PCoA) showed that the community structure of the keystone samples was significantly different from that of no keystone samples (Fig. S8), which implies that keystone taxa may have a significant impact on community structure. Considering that the sample communities with high stability have higher homogeneity under different environmental conditions(40), we further compared the beta diversity of keystone samples and no keystone samples. The results showed that the beta diversity of keystone samples was significantly lower than that of no keystone samples, regardless of whether the abundance and evolutionary position of species were considered in the calculation of distance between samples (Fig. 3b). These results indicated that the community structure variabilities of samples containing keystone taxa were lower than that of samples without keystone taxa. This result provides indirect evidence that the keystone samples have higher community stability than no keystone samples, which was consistent with the results of network analysis.

Furthermore, the alpha diversities of keystone samples, including the Shannon-Wiener index, Pielou’s evenness index, species richness, and phylogenetic diversity, were significantly higher than those of no keystone samples (Fig. S9), which is consistent with another ecological view that high diversity brings high stability(41).

### Samples with keystone taxa have low AS performance in pollutant removal rates

In this study, we measured AS performance using the specific removal rates of different pollutants, including BOD, COD, NH4N, TN, and TP(31). To analyze the impact of keystone taxa on AS performance of AS system, we compared the processing performance between keystone samples and no keystone samples. The results showed that pollutant removal rates of samples containing keystone taxa were significantly lower than those of samples without keystone taxa (Fig. 4a), which indicated that keystone samples had lower performance than no keystone samples. Besides, the comparison of relative abundances of functional groups(42) showed that the low performance of keystone samples may be due to its low abundance of specific functional groups including denitrifiers, polyphosphate-accumulating organisms (PAOs), and glycogen-accumulating organisms (GAOs) (Fig. 4b), not just the large denominator of removal rate caused by the high biodiversity of keystone samples (Fig. S9).

**Fig. 4.**
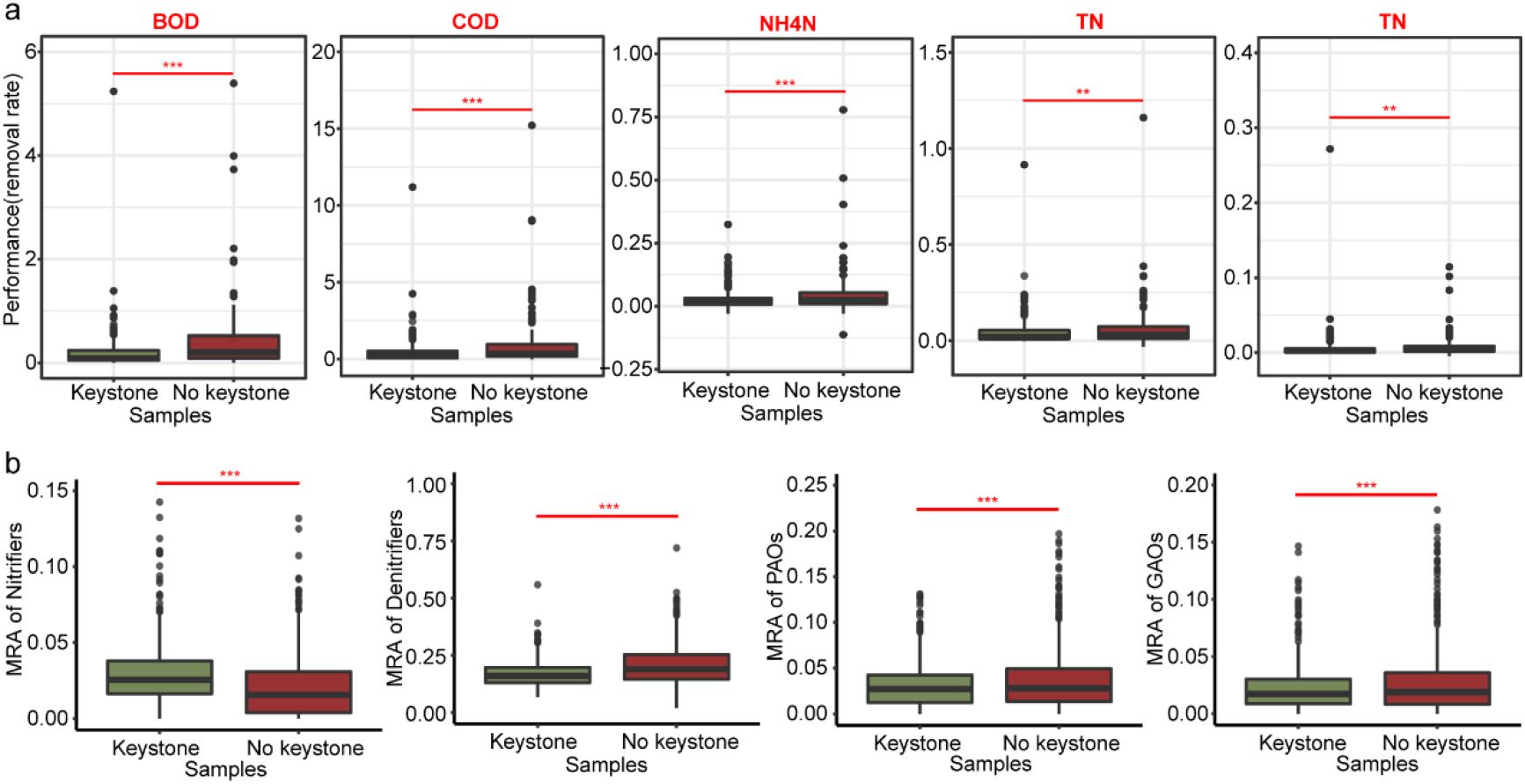
Influence of keystone taxa on AS system functions. **(a)** Comparison of the AS performance between keystone samples and no keystone samples, including BOD, COD, NH4N, TN, and TP removal rate. **(b)** Comparison of the average relative abundance (MRA) of major functional groups between keystone samples and no keystone samples, including nitrifiers, denitrifiers, polyphosphate-accumulating organisms (PAOs), and glycogen-accumulating organisms (GAOs). Statistical analysis was performed using a two-sample Student’s t-test: ***, p < 0.001; **, p < 0.01.

### Correlations between keystone taxa and environmental factors

To investigate key environmental factors significantly related to the keystone taxa, we compared the environmental conditions of keystone and no keystone samples. The quantitative comparison of environmental variables showed that about two-thirds of the environmental factors of the two groups of samples were significantly different (Fig. S10), which may be the reason for the significant differences in the community structure between the keystone samples and the no keystone samples. These significantly different environmental factors include climatic parameters temperature (MAT, AMMaxT, AMMinT, and SMT) and precipitation (SMP), control parameters solid-liquid retention time (HRT, and SRT) and inflow rate (InfR), influx parameters pollutant concentration (InfBOD, InfCOD, InfNH4N, InfTN, and AtInfTP, et. al) and percentage of industrial wastewater (InfPer), and aeration tank parameters sludge volume index (SVI) and sludge load (F/M) (Fig. S10).

To further explore the impact of environmental factors, we took the influx of industrial wastewater as an environmental disturbance, and compared the community structure and AS performance of keystone samples and no keystone samples when the influx water contains or does not contain industrial wastewater. We observed that there were significant differences in the community structures of both keystone samples and no keystone samples when the influx contains industrial wastewater and didn’t contain industrial wastewater (Fig. S11), which indicated that industrial wastewater, as a disturbance factor, significantly affected the microbial community. In addition, we found that the community variability of keystone samples with and without industrial wastewater was significantly lower than that of no keystone samples, suggesting that communities of keystone samples were more stable in response to industrial wastewater disturbance, both in composition and abundance (Fig. 5a).

**Fig. 5.**
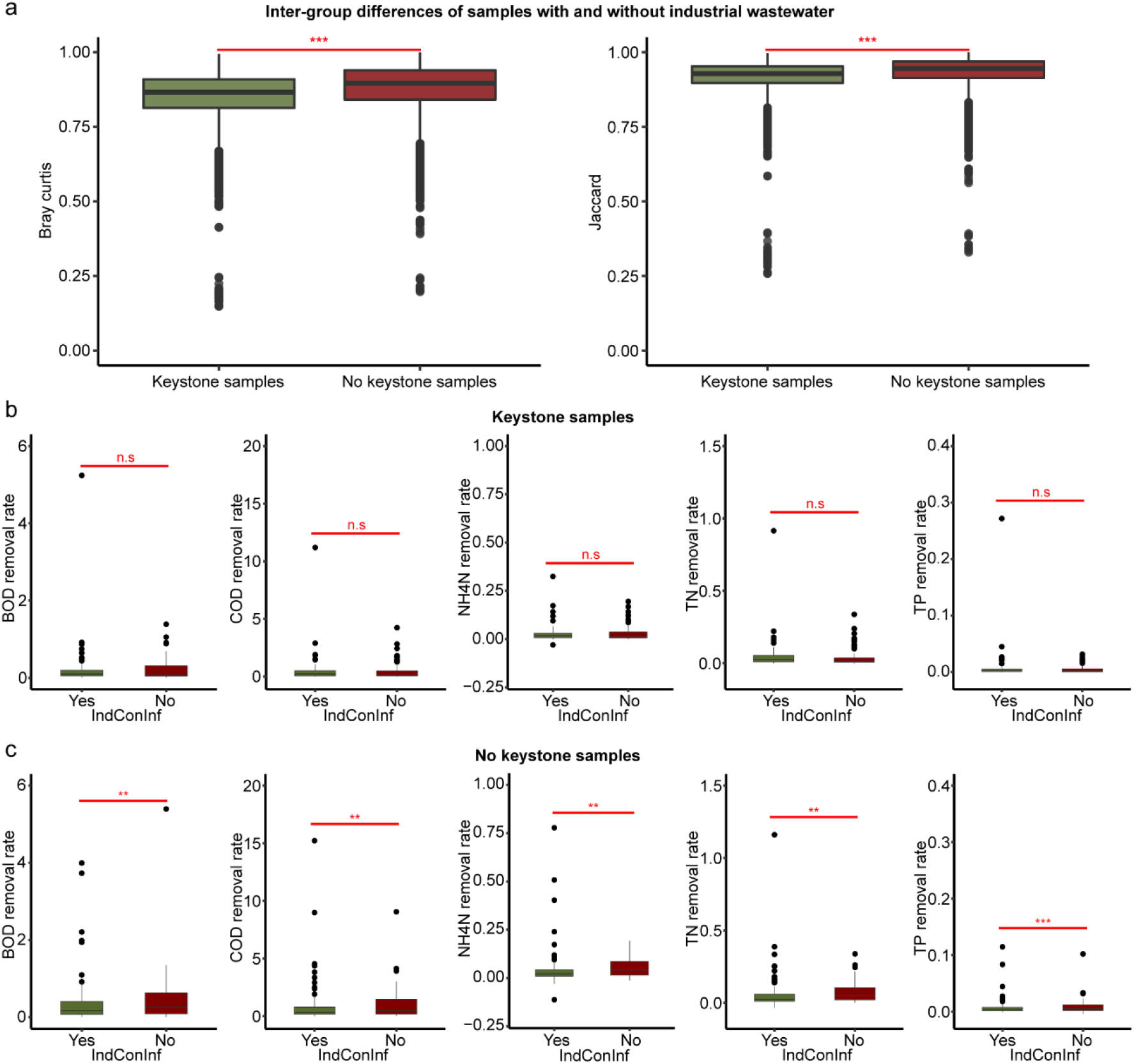
Influence of industrial wastewater in influx on keystone sample and no keystone sample. **(a)** Comparison of inter-group differences of samples with and without industrial wastewater between keystone samples and no keystone samples. **(b)** Comparison of BOD, COD, NH4N, TN, and TP removal rates of the keystone samples with and without industrial wastewater in the influx. **(c)** Comparison of BOD, COD, NH4N, TN, and TP removal rates of the no keystone samples with and without industrial wastewater in the influx. Statistical analysis was performed using a two-sample Student’s *t*-test: ***, p < 0.001; **, p < 0.01; n.s, p > 0.05, no significance.

Furthermore, the comparison results of AS performance show that the shock of industrial wastewater will not affect the community functions of the keystone samples (Fig. 5b), but significantly reduces those of the no keystone samples (Fig. 5c). The result suggested that the communities of the keystone samples show higher resistance to environmental disturbance, which was consistent with their high stability.

### Keystone taxa are more likely to appear in samples with low sludge load

To better understand the causal relationship between environmental factors and the existence of keystone taxa, we calculated the Bayesian network structure between environmental variables and the existence of keystone taxa. The causal relationship derived from the Bayesian network structure showed that the difference in the sludge loading (F/M), the reflux BOD (ReInfBOD), and the sludge volume index (SVI) are key factors that determine the presence of keystone taxa in samples (Fig. 6a).

**Fig. 6.**
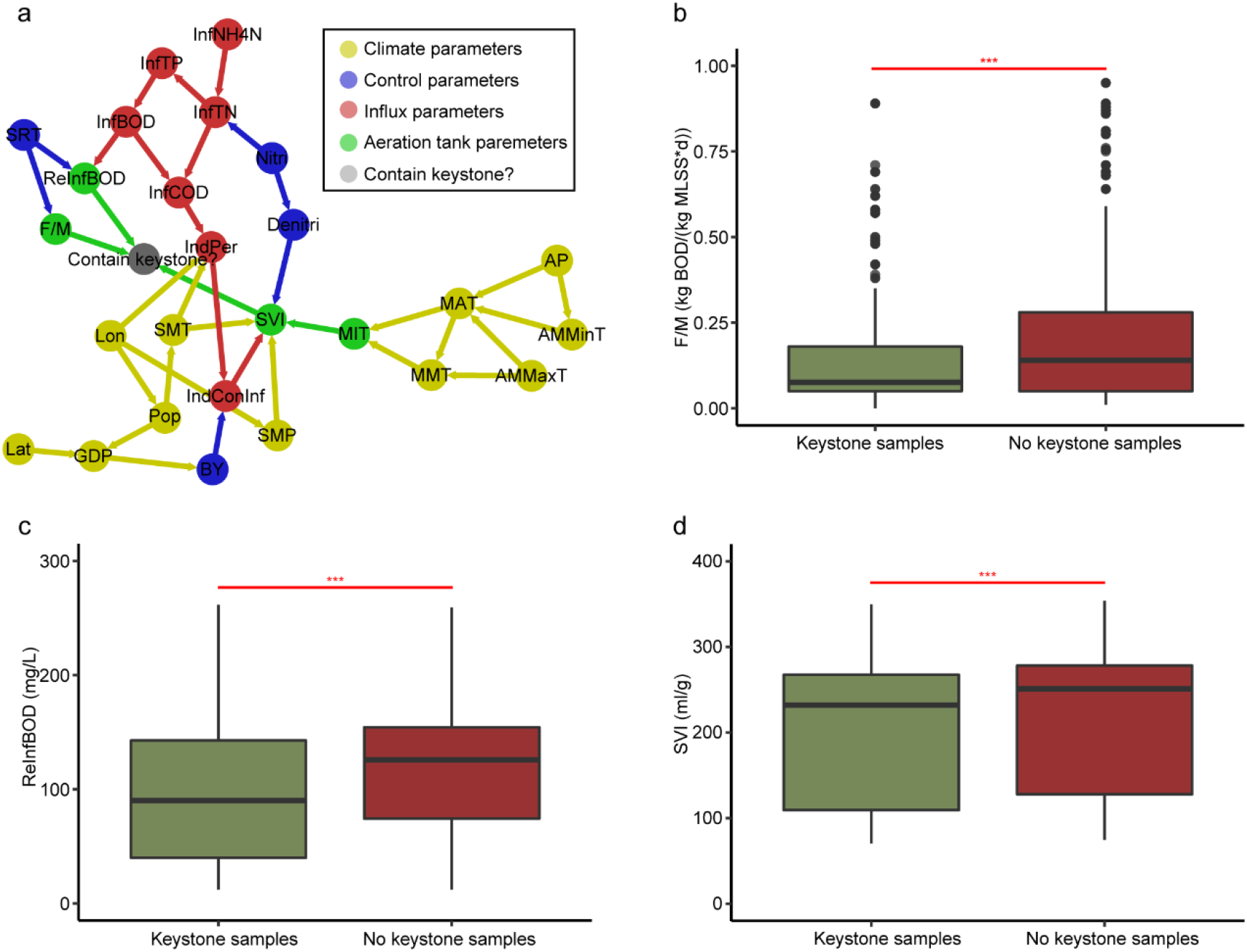
Causal inference of keystone taxa appearing in samples. **(a)** The Bayesian network structure of environmental factors and the occurrence of keystone taxa. Comparison of **(b)** sludge loading, **(c)** reflux BOD and **(d)** sludge volume index between keystone samples and no keystone samples. Statistical analysis was performed using a two-sample Student’s *t*-test: ***, p < 0.001.

We further compared the differences and found that these three factors in the keystone samples were significantly lower than those in the no keystone samples (Fig. 6b; Fig. 6c; Fig. 6d). Previous studies have shown that F/M, ReInfBOD, and SVI are important environmental factors affecting sludge activity(43, 44), and these lower factor values reflect the lower sludge activity to a certain extent. In addition, low sludge activity means low function, so keystone taxa are more likely to appear in samples with low sludge load, low ReInfBOD, and low SVI, which is consistent with the previous conclusion.

Furthermore, the low sludge load of AS system is equivalent to the low nutrient concentration of the ecosystem. Based on the above results, we hypothesize that samples with low nutrient concentration may have the characteristics of high stability and low function of microbial community due to the existence of keystone taxa. To test this hypothesis, the consumer-resource model(45, 46) was used to simulate the microbial community structure under different nutrient concentrations. Here we used average variation degree (AVD) to indicate the community stability(4) and used the resource utilization ratio to reflect the community function. The results showed that with the increase of nutrient concentration, the average variation degree (AVD) of the microbial community increased (Fig. 7a; Spearman’s R=0.835, P<2e-16), and the total resource utilization ratio increased (Fig. 7b; Spearman’s R=0.769, P<2e-16). Therefore, with the increase of nutrient concentration in the sample, the community function will increase while the stability will decrease, which is supported by previous studies(20, 47, 48).

**Fig. 7.**
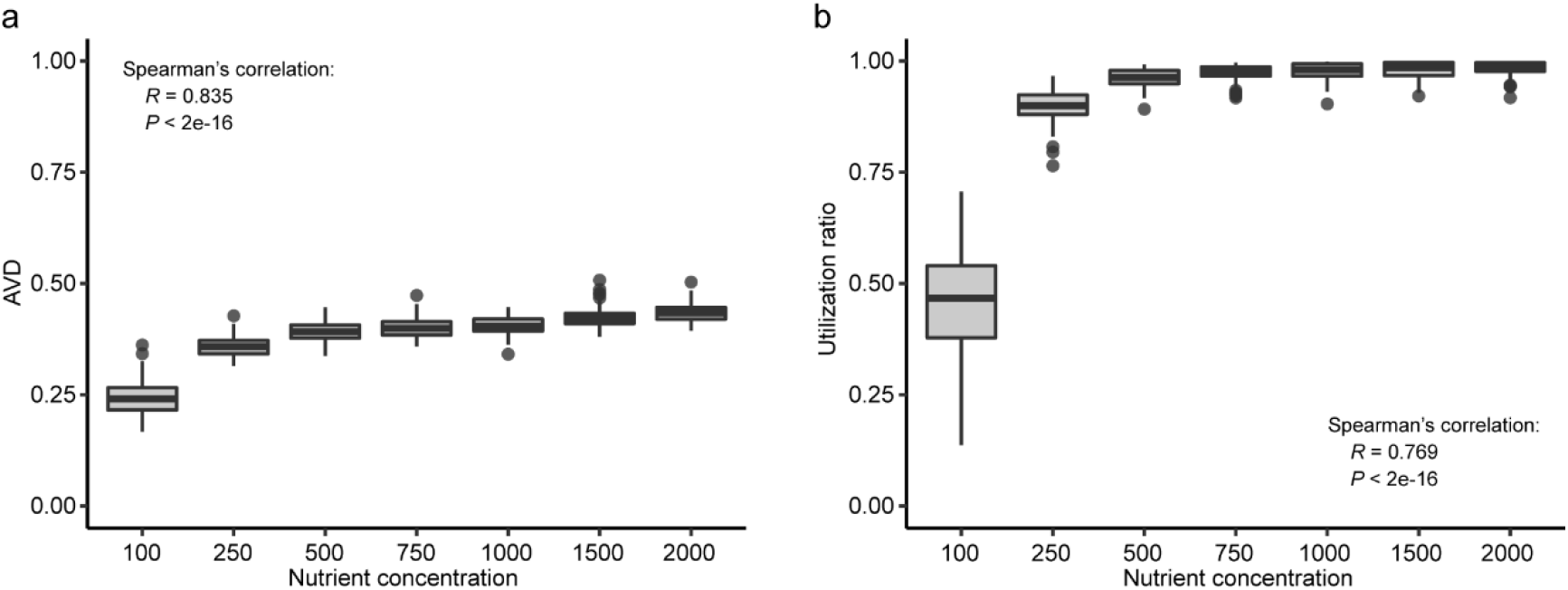
Simulation results of the consumer-resource model. **(a)** The change of average variation degree (AVD) of microbial community with nutrient concentration. **(b)** The change of total resource utilization ratio with nutrient concentration. Spearman correlation analysis is used to describe the correlation between two variables.

## Discussion

In this study, we found AS co-occurrence network is a typical scale-free network, in which a small number of taxa are strongly interconnected. In addition, keystone taxa with high connectivity played an important role in maintaining the microbial community stability of AS system, but at the cost of reducing AS performance. Furthermore, we speculate that keystone taxa are more likely to appear in samples with low sludge load.

Here, the AS network reveals that microbial community assembly in AS system is nonrandom, which is consistent with other ecosystems(24). This indicates that biological interaction plays an important role in community composition and ecosystem function. Furthermore, the keystone defined according to the correlation network well considers the microbial interaction(30, 35), which is quite different from the traditional definition of core species(49). Our results show that keystone taxa defined by this method have a decisive impact on the community stability and function. Further results also suggest that this effect may be caused by the interaction between keystone and other individuals in the community. These results suggest that future research work can try to apply this definition more.

According to the analysis of structural characteristics of keystone taxa, we found that keystone taxa in AS system mainly belonged to rare taxa with low abundance. This shows that keystone taxa emphasize the importance of numerically inconspicuous taxa for community functioning, which is congruent with the rare taxa concept(50). Indeed, the fundamental assumption of keystone taxa and rare taxa is the same: the abundance of a species is not the best determinant of its contribution to the community(30). In addition, keystone taxa emphasized the importance of interaction in the microbial community(35), which is more in-depth than the definition of rare taxa. The importance of rare microorganisms in many biogeochemical processes has been proved(50, 51). Furthermore, due to the abundance disadvantage of rare taxa, they play an important role in the community by providing necessary traits or acting as partners in interspecific interactions(52, 53), and rare taxa act as a reservoir that can quickly respond to environmental changes thus promote community stability(52). Evidence of such low abundant taxa with an over-proportional influence raises the possibility that members of the rare biosphere can also be keystone taxa.

In addition, we noted that in the network stability test of no keystone samples subnetwork, the extinction of the top 30% of species would not significantly affect the natural connectivity of this network (Fig. 3a). Therefore, we compared topology properties of the top 30% species in subnetworks of keystone samples and no keystone samples to analyze the reasons. The comparison results of network topology properties showed that the top 30% of species in the subnetwork of keystone samples had high betweenness centrality and some species also had high node degree (Fig. S12a). While, in the no keystone samples subnetwork, node degrees of the top 30% species were lower, even they also had high betweenness centrality (Fig. S12b). This showed that the node degree of species rather than the betweenness centrality had an impact on the network stability, which is consistent with the previous research results(30, 35).

In the analysis of correlations between the community and environmental factors, we found that the communities harboring keystone taxa had a weaker response to most of the environmental factors, especially the influx factors including industrial wastewater proportion (IndPer) and organic matter content (InfBOD), than the communities of the no keystone samples (Fig. S13). This finding indicates that keystone samples had weak responses to environmental fluctuations derived from substrate influx. On the one hand, Industrial wastewater contains toxic and harmful substances(54, 55). Therefore, IndPer reflects the interference of toxic and harmful substances to AS system. The small correlation with IndPer would lead to higher community stability and lower community variability of keystone samples (Fig. 3b). On the other hand, InfBOD reflects the organic load of AS system. The small correlation with InfBOD indicated that keystone samples were less flexible to the organic matter in the influx, which made the community unable to adjust their structure in response to the pollutants influx, so their removal rates would be lower (Fig. 4a). Therefore, the existence of keystone taxa is a double-edged sword. It maintains the stability of the community when the environment is not suitable for survival, but also reduces the efficiency of sewage treatment when the influent load is large.

Furthermore, we found that the keystone taxa often appeared in samples with low sludge load, which had high community stability and low function. High sludge load imposes higher substrate toxicity and pressures on the microbial communities, which drives the communities to respond with increasing removal performance to reduce these pressures(56). The significant positive correlations between sludge load and pollutant degradation rates support this idea (Fig. S14). Furthermore, we used the consumer-resource model to verify that nutrient concentration has opposite effects on community stability and function, which is supported by several literatures(20, 47, 48). Some studies have found that stability is not conducive to function(57). However, the role of keystone taxa in this process needs more experimental evidence to prove.

Previous studies have shown that definitions of stability in ecology can be classified generally into two categories, one is based on a system’s dynamic stability, and the other is based on a system’s ability to defy change (resilience and resistance)(14). In this paper, the natural connectivity of AS network under simulated species extinction reflects the community stability in terms of resistance(5), and the beta diversity of the community reflects the community variability of different samples, which to some extent reflects the stability of the system dynamics(58). However, the variability of community structure, on the one hand, reflects the community instability, on the other hand, may also be due to the difference in environmental conditions (Fig. S15a). To control the impact of environmental differences, we divided the beta distance between samples by the environmental distance and get the corrected beta distance to reflect the community instability. The results indicate that the keystone sample community was more stable than no keystone samples (Fig. S15). Nevertheless, if there are no significant differences under other environmental conditions, but only differences in whether to contain keystone taxa, the demonstration of the system’s dynamic stability here will be more rigorous.

In summary, our results demonstrate that keystone taxa can help identify stable communities like a biomarker and explore the relationship between community stability and function. These findings provide new insights into the understanding of the relationships between microbial community stability and function and highlight the important role of keystone taxa in the ecosystem.

## Methods

### Datasets and data preprocessing

This study used a previously published dataset of 1186 activated sludge samples taken from 269 WWTPs in 23 countries across 6 continents(31). In addition to 16S rRNA sequencing data of these sludge samples, associated metadata conforming to the Genomic Standards Consortium’s MIxS and Environmental Ontology Standards(59) were also provided by plant managers and investigators.

The original microbial sequencing data were processed using Quantitative Insights Into Microbial Ecology (QIIME2) software (http://qiime2.org)(60). All paired reads were merged, quality filtered, then denoised through the DADA2 plugin(61) to clustered into 100% amplicon sequence variants (ASVs). Then, ASVs classified as fungi, ASVs with unassigned taxonomy at the domain level, and ASVs annotated as mitochondria or chloroplasts were removed so that only bacterial and archaeal sequences were retained. Singletons (ASVs with only one sequence) were discarded before further analysis to reduce the impact of sequencing errors. Then we rarefied each sample to 20954 sequences, to obtain the maximal observation of both samples and features, from which 51998 ASVs were obtained. The final feature sequences were taxonomically and functionally classified using the MiDAS 4 reference database(42).

To have a clearer understanding of the metadata, we classified environmental factors into different types(31), including climate parameters, control parameters, influx parameters, and aeration tank parameters of samples (Table S2). To reduce the redundancy of metadata, we first removed some non-numerical variables of multiple categories that are difficult to operate and some variables with no practical meanings, such as site name, city name, etc. Then, the LabelEncoder algorithm was used to numeric binary non-numeric variables, and missing values were completed according to the two-nearest neighbor principle (Table S3).

### Co-occurrence network of AS system

To reduce noise and false-positive predictions, network inclusion was restricted to ASVs with average relative abundance greater than 0.001% and present in at least 1% of samples (1, 2), comprising 6,752 ASVs in total. Then, the microbial co-occurrence network of the global AS system was constructed based on Pearson correlations of log-transformed ASV abundances(6), followed by a random matrix theory (RMT) based approach that automatically determines the correlation cut-off threshold (62). The final AS network contains 4992 nodes (Table S4) and 65457 edges.

Network visualization was generated with Gephi (v. 0.9.7). In the AS network, each node indicates a given ASV, and each edge represents a significant correlation between two ASVs. The degree represents the number of edges connecting each node to the rest nodes of the network. Normally, the high topological characteristic values (such as node, edge, and degree) suggest a more complex network. In addition, 100 Erdös -Réyni random networks, which exhibit the same number of nodes and edges as the AS network, were calculated using the igraph package of R (v. 4.0.3), with each edge having the same probability of being assigned to a node(63). To further describe the topological parameters of the network, a set of metrics of both AS and random networks were calculated and compared, including clustering coefficient, average path length, and modularity.

Furthermore, node centrality measures are tools designed to identify node importance(64). However, node importance is a rather vague concept and can be interpreted in various ways, giving rise to multiple coexisting centrality measures, the most common being degree, betweenness, closeness, and eigenvector centrality. In degree centrality, the importance of a node is measured by the number of nodes it can immediately influence. In betweenness centrality, the importance of a node is given by the frequency of this node belonging to the shortest path between the other two nodes in the network. In closeness centrality, importance is measured in terms of how fast information can travel from a given node to every other node in the network. In eigenvector centrality, the importance of a node is computed as a function of the importance of its neighbors. All these node centralities were calculated using the igraph and tnet package of R and were shown in Table S5.

Finally, network stability was evaluated by removing nodes in the AS network to estimate how quickly robustness degraded, and network robustness was assessed by the natural connectivity of the nodes(65).

### Defining keystone taxa

Recent studies have shown that microbial communities harbor keystone taxa, which drive community composition and function irrespective of their abundance(30). Network analysis can be a powerful tool for inferring keystone taxa from microbial communities. Although high betweenness centrality was previously used to identify keystone taxa statistically in several studies(23, 66), it was recently shown that high mean degree, high closeness centrality, and low betweenness centrality can be collectively used to identify keystone taxa with 85% accuracy(35). In this study, nodes with high degrees (>100) and low betweenness centrality values (<5000) are recognized as keystone taxa of AS system(24).

### Defining hubs and core taxa

Based on metabolic network approaches, the network hubs (Zi > 2.5, Pi > 0.62; i.e., “Zi” indicates within-module connectivities and “Pi” indicates among-module connectivities), module hubs (Zi > 2.5, Pi < 0.62), connectors (Zi < 2.5, Pi > 0.62), and peripherals (Zi < 2.5, Pi < 0.62) were identified(28). All network hubs, module hubs, and connectors were highly linked to their own or others’ modules, which were called hubs taxa of AS system.

The global-scale core taxa of WWTP was determined based on multiple reported measures(31). First, ‘overall abundant ASVs’ were filtered out according to the mean relative abundance across all samples. We selected all top 1% ASVs as overall abundant ASVs. Second, ‘ubiquitous ASVs’ were defined as ASVs with an occurrence frequency in more than 20% of all samples. Finally, ‘frequently abundant ASVs’ were selected based on their relative abundances within a sample. In each sample, the ASVs were defined as abundant when they had a higher relative abundance than other taxa and made up the top 80% of the reads in the sample(67). A frequently abundant ASV was defined as abundant in at least 10% of the samples. Following the same criteria described above, a core ASV should be one that was from the top 1% ASVs, a core ASV also had to be detected in more than 20% of the samples and dominant for more than 10% of the samples. Corresponding to the core taxa, ASVs that did not meet the above three conditions were called non-core taxa.

### Subnetwork of keystone samples and no keystone samples

The sample subnetwork is a subset of the co-occurrence network, which contains the species in the sample and all its related edges. We generated subnetworks for two sample groups from the AS co-occurrence network by preserving ASVs presented in each sample group and their related edges using *subgraph* functions in igraph packages. The topological properties of these subnetworks were also calculated using the igraph package of R (v. 4.0.3).

### Bayesian network structure

Bayesian networks (BNs) are a type of probabilistic graphical model consisting of a directed acyclic graph. In a BN model, the nodes correspond to random variables, and the directed edges correspond to potential conditional dependencies between them. The Bayesian network structure is the relationship framework between nodes of BNs. Obtaining the Bayesian network structure can obtain the qualitative relationship between variable nodes.

Previous theoretical research showed that Bayesian network structures can be obtained in two ways, one is based on statistical analysis, and the other is based on scoring search. Here, we used the *h2pc* function in the bnlearn package of R to obtain the Bayesian network structure between environmental factors. H2pc is a hybrid algorithm, which greatly improves the computational efficiency of the algorithm based on ensuring the accuracy of the results.

### Consumer-resource model

The microbial consumer-resource model is the smallest model of a microbial community growing in an environment with limited mixed resources(45, 46), which simulates species and resources change with time and usually is used to describe the community dynamics. The implementation of this model is generated using an open-source Python package Community Simulator(68). In this study, the dynamic process of the microbial community under different nutrient concentrations (100, 250, 500, 750, 1000, 1500, 2000) was simulated, and each concentration condition was paralleled 100 times. For the microbial community structure under different simulated steady-state conditions, the average variation degree (AVD) was calculated to reflect the community stability(4). In addition, the resource utilization ratio of the community at the end of the simulation was calculated to reflect the community function.

### Statistical analysis

All alpha diversity measures were conducted using the vegan and Picante packages of R (v. 4.0.3). Unless indicated otherwise, an unpaired, two-tailed, two-sample Student’s *t*-test was performed for comparative statistics using the *t*.*test* function in the stats package. Non-linear fitting was implemented using the *nls* function in the stats package. The statistical comparison of slopes was realized by using the *diffslope* function in the simba package. All analysis and graphing were done using R 4.0.3 or python 3.8.

## Acknowledgments

The authors thank the Global Water Microbiome Consortium (GWMC) and all the people involved for providing samples and plant metadata. We thank the High-performance Computing Platform of Peking University for providing the computing platform.

## Author’ contributions

All authors contributed intellectual input and assistance to this study. X.L. conceived the study and performed the original analysis. B.L. added simulation validation. X.L. and Y.N. co-wrote the paper. M.W., X.C., L.A., Y.N., and X.L.W. revised it. All authors discussed the results and commented on the article.

## Availability of data and materials

The raw data in this study is from reference 31. The source code is available at https://github.com/Neina-0830/WWTP_community_network.

## Ethics approval and consent to participate

Not applicable.

## Consent for publication

Not applicable.

