## SUPPLEMENTARY INFORMATION for "Keystone taxa responsible for the microbial community stability and performance of activated sludges"

#### For

**This PDF file includes:**

Supplementary Figs. 1 to 15

Supplementary Tables 1 to 5

### Supplementary Figures

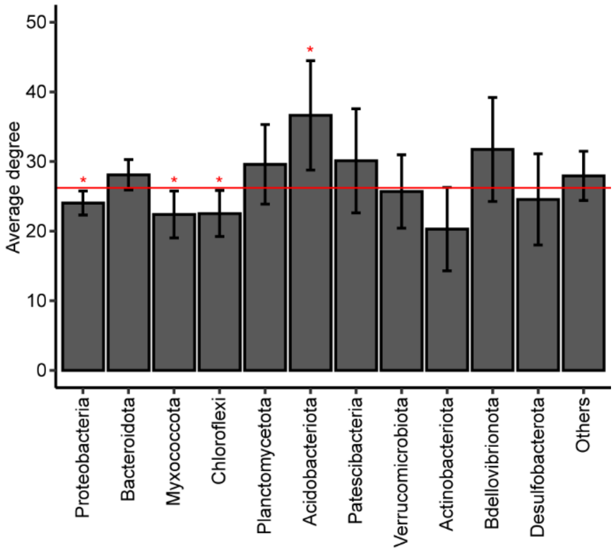


**Fig. S1 Average node degree of different phyla in the microbial co-occurrence network of AS system.** The error bar represents the upper and lower 95% confidence intervals of the mean value. The significance shows the comparison between node degrees of different phyla and the average node degree of AS network. Statistical analysis was performed using a two-sample Student's t-test: ***, $p < 0.001$.

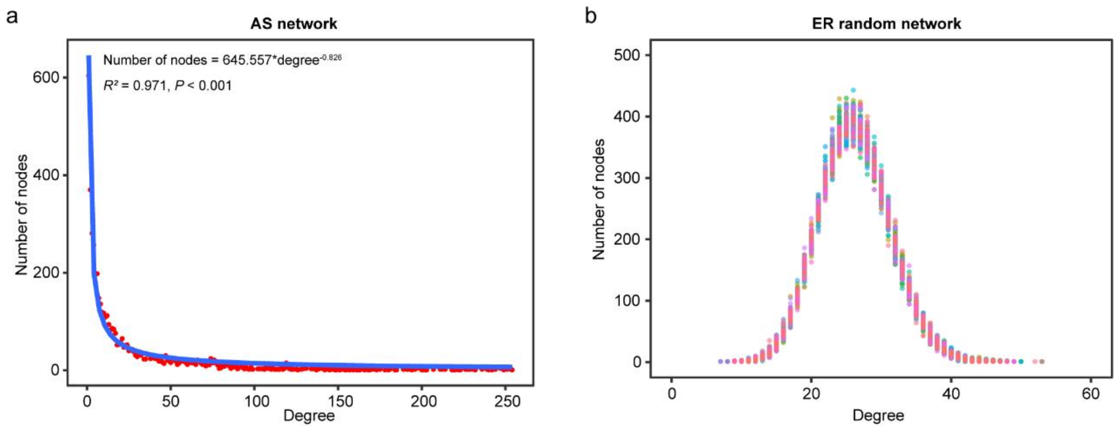


**Fig. S2 Node degree distribution of co occurrence network. (a)** Power-law distribution of node degree of AS network. **(b)** Poisson distribution of node degree of ER random network.

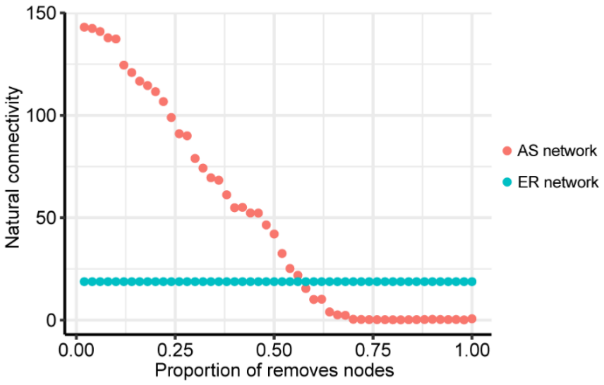


**Fig. S3 Stability of AS network.** Stability is measured by the natural connectivity of the network.

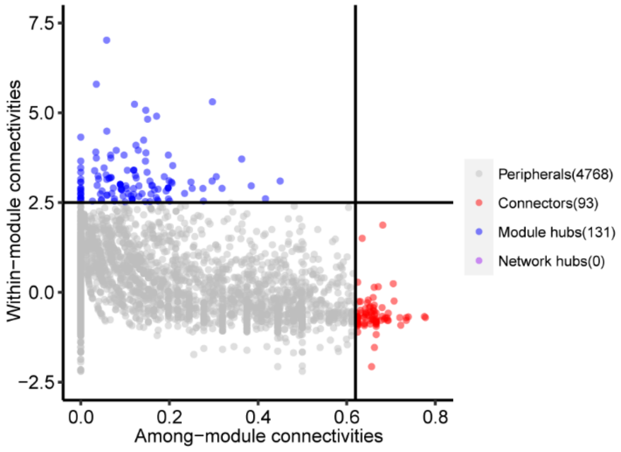


**Fig. S4 Network hubs in AS network.** Network hubs are divided by within-module (Zi) and among-module (Pi) connectivity. Where, peripheral nodes ($Zi \leq 2.5$, $Pi \leq 0.62$), connectors ($Zi \leq 2.5$, $Pi > 0.62$), module hubs ($Zi > 2.5$, $Pi \leq 0.62$), and network hubs ($Zi > 2.5$, $Pi > 0.62$). We refered to connectors, module hubs and network hubs as the hubs in the network.

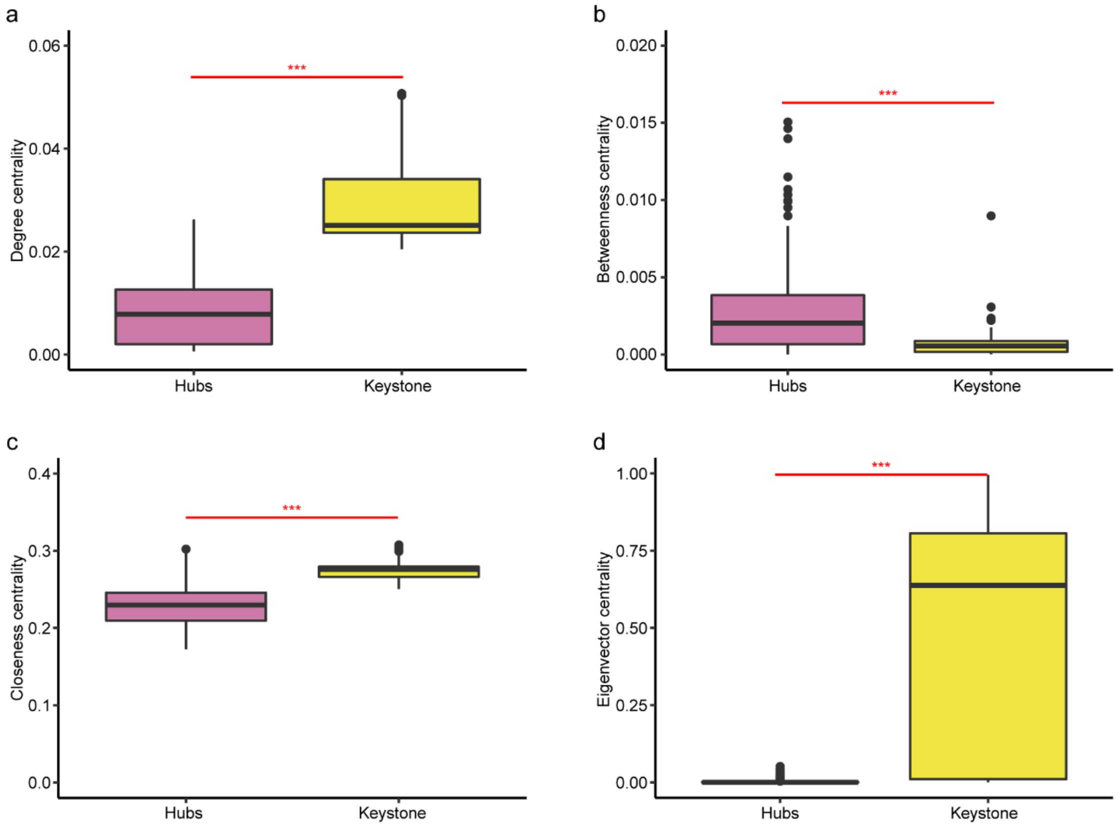


**Fig. S5 Comparison of centrality between hubs and keystone taxa.** Comparison of **(a)** degree centrality, **(b)** betweenness centrality, **(c)** closeness centrality and **(d)** eigenvector centrality between keystone taxa and other taxa. Statistical analysis was performed using a two-sample Student's t-test: ***, $p < 0.001$.

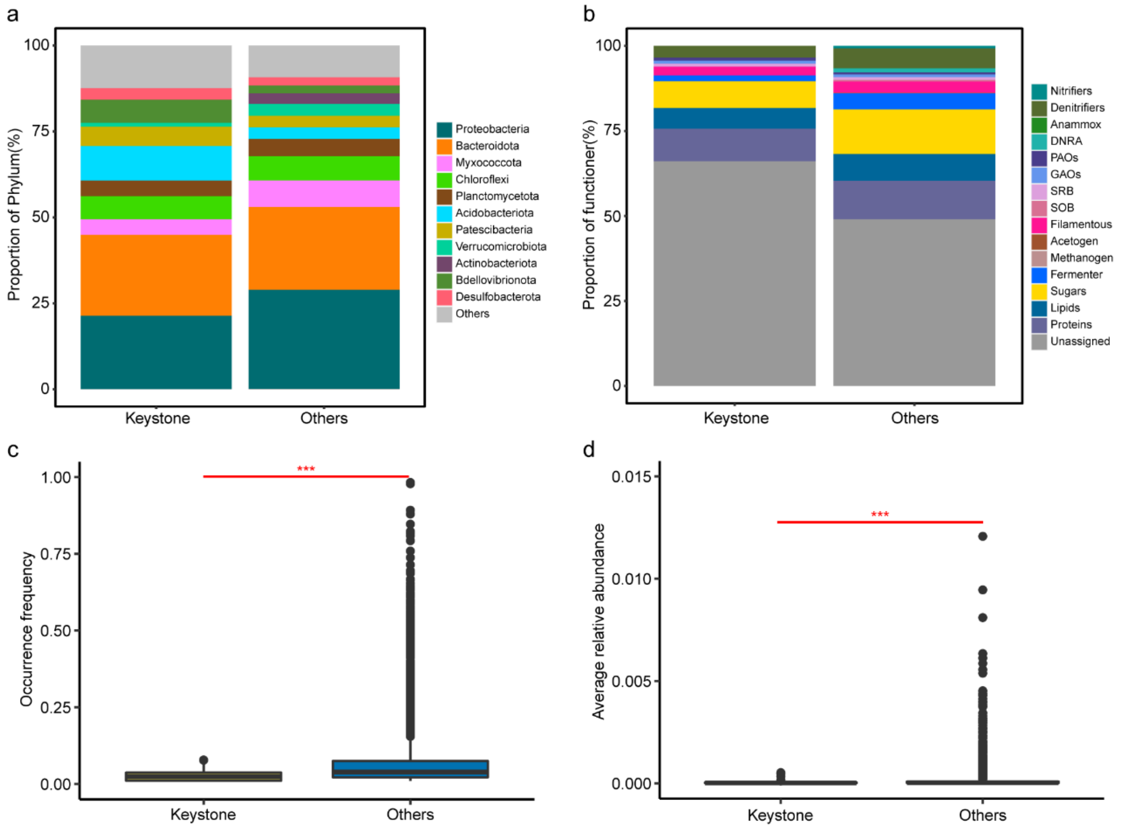


**Fig. S6 Comparison of community structure characteristics between keystone taxa and others taxa.** Comparison of **(a)** phylum and **(b)** functional composition between keystone taxa and other taxa. Comparison of **(c)** occurrence frequency, **(d)** average relative abundance between keystone taxa and other taxa. Statistical analysis was performed using a two-sample Student's t-test: ***, $p < 0.001$.

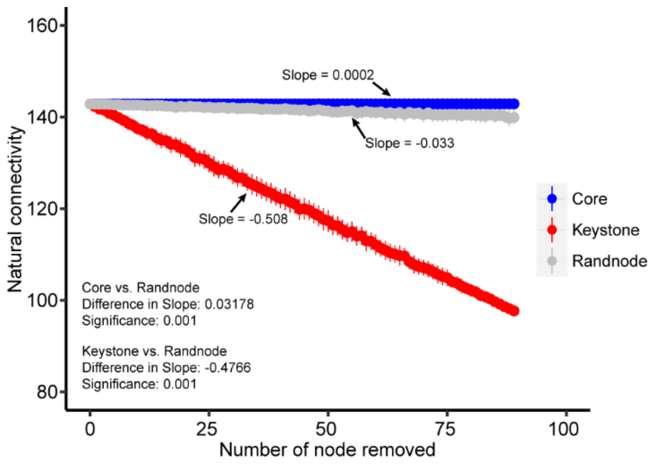


**Fig. S7 Comparison of core taxa and keystone taxa impact on AS network stability.** The impact of randomly deleting keystone taxa or core taxa on the natural connectivity of AS network. The statistical comparison of slopes was realized by using *diffslope* function in simba package of R 4.0.3.

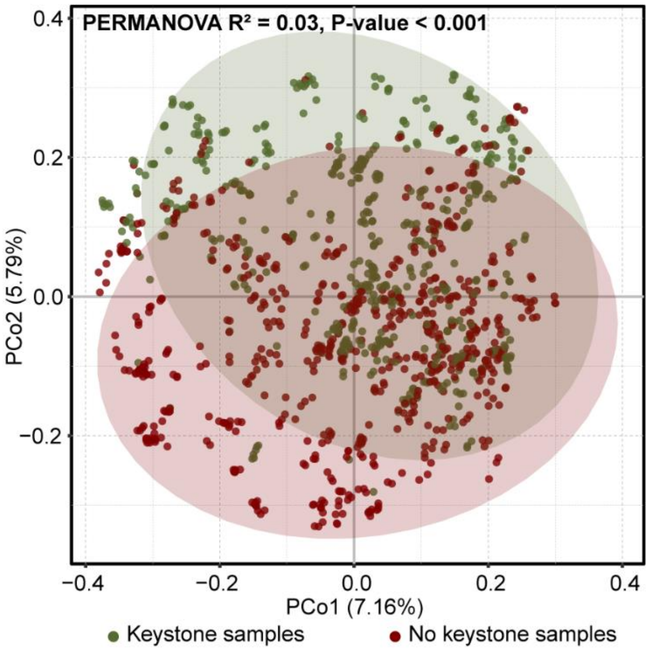


**Fig. S8 Comparison of community structure between keystone samples and no keystone samples.** The principal coordinate analysis (PCoA) analysis is based on ASVs table and the difference significance analysis is based on PERMANOVA.

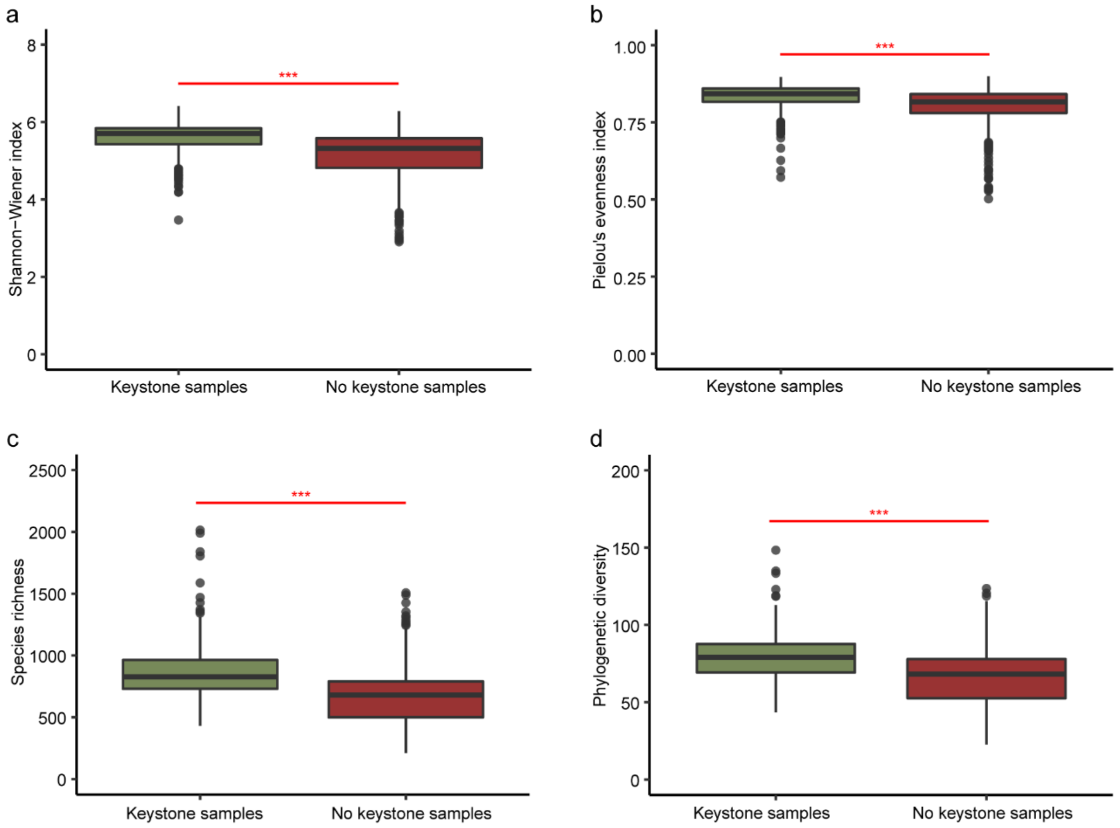


**Fig. S9 Comparison of alpha diversity indexes between keystone samples and no keystone samples.** Comparison of **(a)** Shannon-Wiener index, **(b)** Pielou's evenness index, **(c)** species richness and **(d)** phylogenetic diversity of the two types of samples. Statistical analysis was performed using a two-sample Student's t-test: ***, $p < 0.001$.

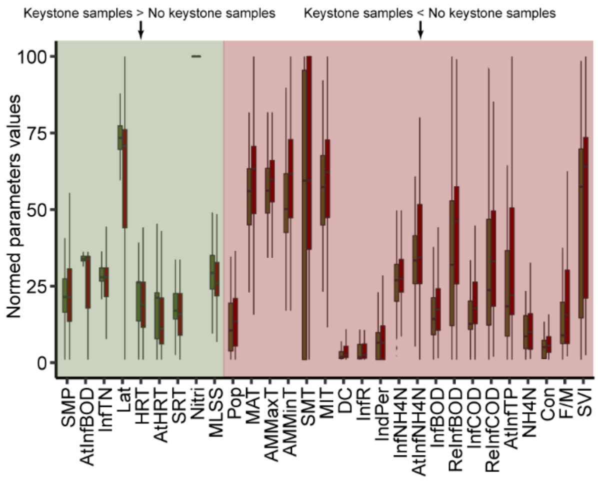


**Fig. S10 Environmental factors with significant difference between keystone samples and no keystone samples.** Light green shows that the environment factor of keystone samples is higher than that of no keystone samples. Light red shows that the environment factor of keystone samples is lower than that of no keystone samples. Statistical analysis was performed using a two-sample Student's *t*-test, and the p value of environmental factors with significant difference is lower than 0.05.

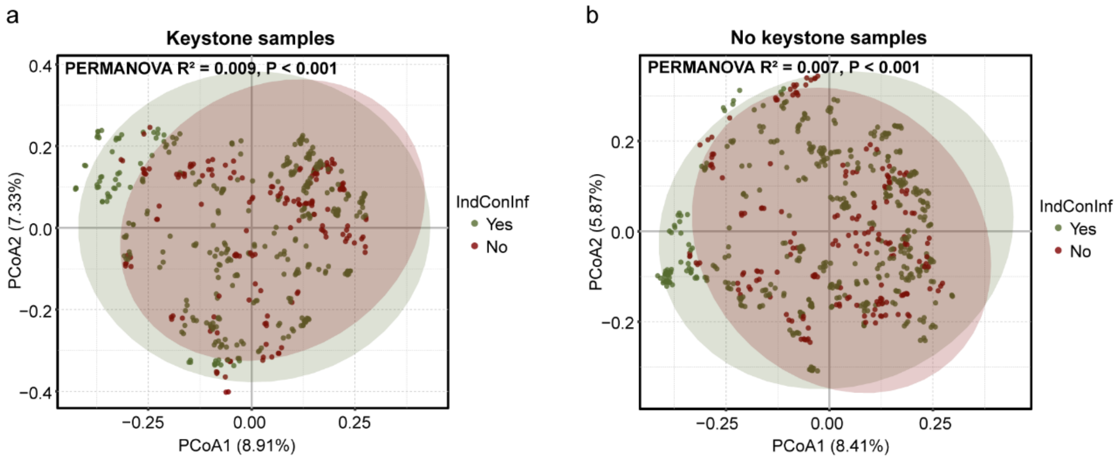


**Fig. S11 Comparison of the community structure when the influx water contains or does not contain industrial wastewater. (a)** Comparison of the community structure of keystone samples when the influx water contains or does not contain industrial wastewater. **(b)** Comparison of the community structure of no keystone samples when the influx water contains or does not contain industrial wastewater. The principal coordinate analysis (PCoA) analysis is based on ASVs table and the difference significance analysis is based on PERMANOVA.

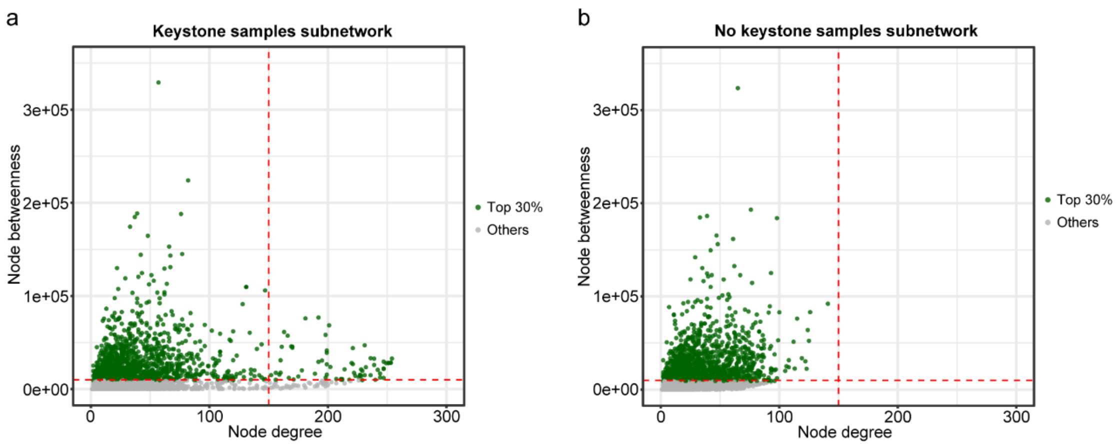


**Fig. S12 Topological properties of the top 30% species in subnetworks of keystone samples and no keystone samples. (a)** Topological properties of the top 30% species in keystone samples subnetwork. **(b)** Topological properties of the top 30% species in no keystone samples subnetwork.

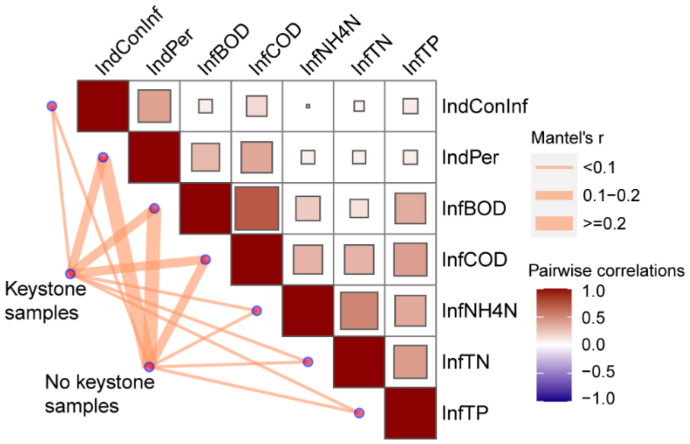


**Fig. S13 Correlation analysis of influx conditions with community structure of keytsone samples and no keystone samples.** Edge width corresponds to the Mantel's r value. Pairwise correlations of these factors are shown with a color gradient.

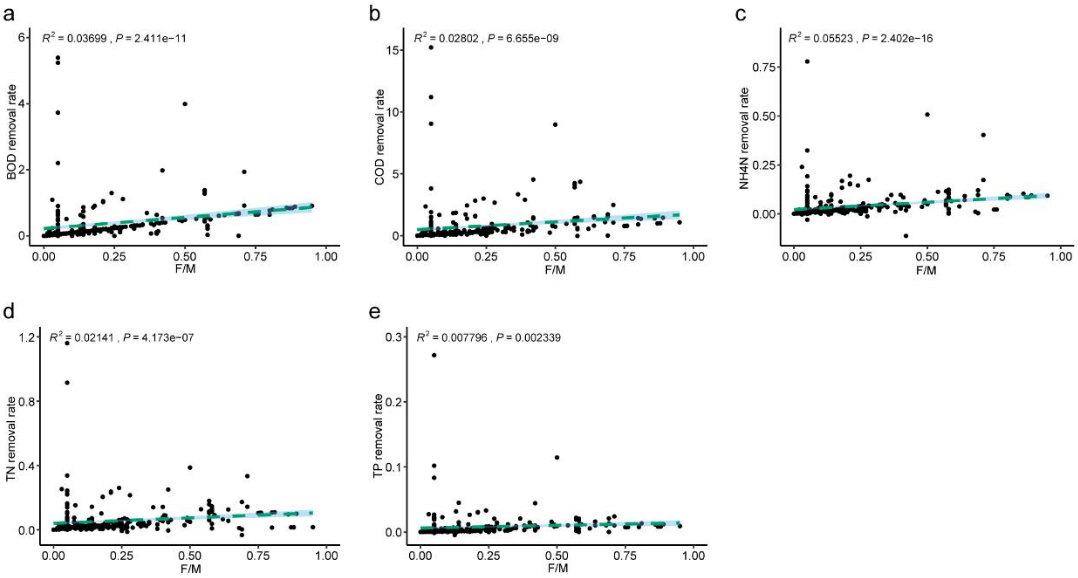


**Fig. S14 Linear correlation between sludge load and pollutant removal rate.** Linear regression results between sludge load and **(a)** BOD, **(b)** COD, **(c)** NH4N, **(d)** TN, and **(e)** TP removal rate. The $R^2$ and P values were goodness of fit and significance of linear regression respectively.

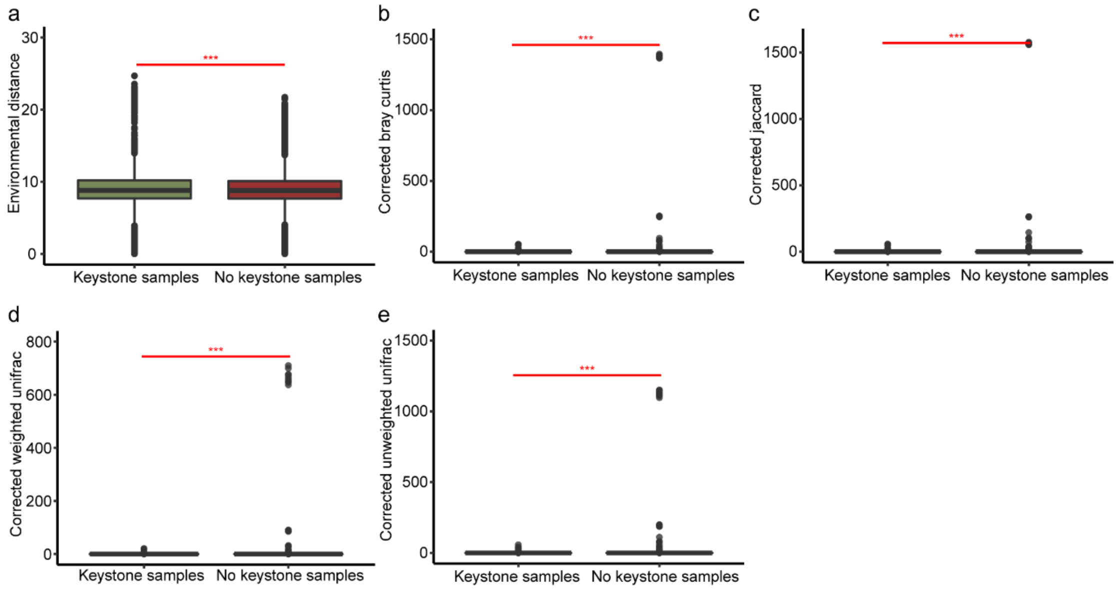


**Fig. S15 Comparison of environmental and community variability between keystone samples and no keystone samples. (a)** Comparison of environmental variability between keystone samples and no keystone samples based on euclidean distance. Comparison of corrected community variability between keystone samples and no keystone samples based on **(b)** bray curtis, **(c)** jaccard, **(d)** weighted unifrac, and **(e)** unweighted unifrac distance.

### Supplementary Tables

**Supplementary Table 1. Topological properties of keystone samples subnetwork and no keystone samples subnetwork.**

| | node | edge | Average degree (AD) | Clustering coefficient (CC) | Average path length (APL) | Modularity (MD) |
|---|---|---|---|---|---|---|
| Keystone samples subnetwork | 4811 | 58991 | 24.523 | 0.581 | 5.090 | 0.771 |
| No keystone samples subnetwork | 4746 | 44729 | 18.849 | 0.507 | 5.133 | 0.820 |

**Supplementary Table 2. Abbreviations and meanings of environment factors.**

This table is available as a supplementary dataset; Table S1.xlsx.

**Supplementary Table 3. Numerical and normalized environmental data.**

This table is available as a supplementary dataset; Table S2.xlsx.

**Supplementary Table 4. Basic information of nodes in AS co-occurrence network.**

This table is available as a supplementary dataset; Table S3.xlsx.

**Supplementary Table 5. Centralities of nodes in AS co-occurrence network.**

This table is available as a supplementary dataset; Table S4.xlsx.
